# MCT1 governs a metabolic checkpoint at pachytene during spermatogenesis

**DOI:** 10.64898/2026.04.21.720010

**Authors:** Xiaoyu Zhang, Yan Liu, Ning Wang

## Abstract

The transition from mitosis to meiosis represents a fundamental cell-fate decision that requires coordinated remodeling of transcriptional and metabolic programs. While key transcriptional regulators of meiotic entry have been defined, how metabolic flux directly governs this process remains unclear. Here, we identify a monocarboxylate transporter1 (MCT1)-dependent metabolic checkpoint that controls meiotic progression in mammalian spermatogenesis. Through integrative single-cell transcriptomics, metabolic profiling, and computational perturbation modeling, we show that *Stra8*-driven meiotic initiation is coupled to a metabolic switch favoring monocarboxylic acid metabolism, prominently involving MCT1 (encoded by *Slc16a1*). Germ cell–specific deletion of *Slc16a1* results in a complete arrest at the pachytene stage, characterized by defective homologous recombination, persistent DNA damage, and failure to activate the meiotic transcriptional program. Multi-omic analyses reveal that loss of MCT1 induces a metabolic stress-like state, suppresses expression of key meiotic regulators, and disrupts progression through the pachytene checkpoint. Mechanistically, we demonstrate that MCT1-mediated lactate influx drives histone H4 lysine 12 lactylation (H4K12la) at promoters of meiotic genes, thereby epigenetically licensing their expression. In the absence of MCT1, H4K12la deposition is lost at meiotic loci and redistributed toward stress-response pathways. Together, our findings suggest MCT1-mediated metabolism as an instructive signal that integrates metabolic state with epigenetic regulation to govern meiotic cell-fate progression, defining a previously unrecognized metabolic checkpoint at pachytene.

## INTRODUCTION

The transition from mitosis to meiosis is a defining event in the life cycle of all sexually reproducing organisms. In mouse germ cells of both sexes, this process is regulated by retinoic acid (RA) through its target, stimulated by retinoic acid gene 8 (Stra8)(Ref. 1-3). *Stra8*-deficient germ cells fail to enter meiosis^2^. Together with its cofactor MEIOSIN, Stra8 regulates meiotic gene expression by binding to meiotic gene loci^4,5^; however, neither Stra8 nor MEIOSIN is sufficient to activate their transcription^4,6^. Single-cell RNA sequencing analysis further shows that *Stra8*-deficient germ cells can progress to a leptotene-like stage, where they exhibit transcriptional features resembling leptotene spermatocytes but lack meiotic gene expression (e.g., *Dmc1, Hormad1, Sycp3*) (ref. 7).

Upon entry into meiosis, germ cells initiate meiotic prophase I, a highly orchestrated program that ensures accurate homologous chromosome segregation and genetic diversity through recombination^8^. Meiotic prophase I is subdivided into four sequential stages—leptotene, zygotene, pachytene, and diplotene—each characterized by distinct chromosomal and molecular events. During leptotene, chromosomes begin to condense and programmed DNA double-strand breaks (DSBs) are introduced by the Spo11 complex to initiate recombination. In zygotene, homologous chromosomes undergo pairing and synapsis, mediated by assembly of the synaptonemal complex.

Pachytene represents a critical stage at which synapsis is complete and homologous recombination proceeds, allowing crossover formation and genetic exchange. At this stage, cells are subject to the pachytene checkpoint—the most stringent of the meiotic checkpoints—which monitors the completion of homologous synapsis and the repair of programmed DNA DSBs; defects in these processes trigger cell cycle arrest and apoptosis^9^. Finally, during diplotene, the synaptonemal complex disassembles, homologous chromosomes begin to separate while remaining connected at chiasmata, and cells prepare for subsequent meiotic divisions.

These meiosis-specific chromosomal remodeling events requires precise activation of a specialized gene regulatory network distinct from that of mitosis^5,7,10^. While past studies have identified key transcription factors (e.g., Stra8, MEIOSIN) orchestrating meiosis, a fundamental question remains: how do germ cells meet the metabolic and regulatory decision associated with this transition? Traditionally, metabolic reprogramming has been viewed primarily as a supportive process to satisfy adenosine triphosphate (ATP) requirements. However, accumulating evidence in stem cell biology suggests that metabolic states and specific metabolites can actively influence cell fate decisions and progenitor behavior^11,12^. Whether metabolic flux directly governs the gene regulatory networks required for meiotic commitment remains largely unknown.

Notably, in mammalian spermatogenesis, the interplay between metabolism and cellular differentiation is particularly pronounced. Accompanying the transition from mitosis to meiosis, male germ cells traverse the blood-testis barrier (BTB) (ref. 13). As a result, meiotic germ cells are isolated from the circulation and exhibit a distinct metabolic profile. According to the classical lactate shuttle paradigm, meiotic germ cells rely on lactate produced by Sertoli cells via anaerobic glycolysis, which metabolize glucose and release lactate into the seminiferous environment, rather than directly utilizing circulating glucose^14,15^. In addition to serving as an energy substrate, lactate also supports germ cell survival^16^. Interestingly, recent studies have revealed that lactate functions not only as a metabolic substrate but also as a regulator of epigenetic modifications through histone lactylation^17^. In spermatogenesis, histone lactylation is enriched not only at meiotic gene loci but also at recombination hotspots, suggesting a broader role for lactate beyond metabolism^18^. However, whether lactate-dependent metabolic flux directly governs meiotic progression, and how such metabolic signals are integrated into the gene regulatory network controlling meiosis, remain unresolved.

Here, we address this fundamental question by investigating how metabolic state influences the mitosis-to-meiosis transition and subsequent meiotic progression. Using integrative single-cell transcriptomics, computational modeling, and genetic approaches, we identify a Stra8-dependent metabolic switch that promotes monocarboxylic acid metabolism and highlights a central role for the lactate transporter MCT1 (encoded by *Slc16a1*). We demonstrate that germ cell–specific loss of MCT1 results in a stringent arrest at the pachytene stage, characterized by defective homologous recombination and failure to activate the meiotic transcriptional program. Mechanistically, we find that MCT1 is required for histone H4 lysine 12 lactylation (H4K12la) at meiotic gene promoters, linking monocarboxylate transport to epigenetic regulation of meiotic gene expression. Together, our findings reveal a previously unrecognized metabolic checkpoint at pachytene and establish lactate metabolism as an instructive regulator that couples metabolic flux to epigenetic control of meiotic cell fate.

## RESULTS

### Stra8 drives a monocarboxylic acid metabolic program during meiotic initiation

To delineate the transcriptional metabolic landscape underlying the Stra8-mediated transition from mitosis to meiosis, we reanalyzed our previously published single-cell RNA sequencing (scRNA-seq) dataset comprising wild-type and *Stra8*-deficient juvenile testicular cells^7^. Focusing specifically on metabolic gene signatures (1,731 genes) within the early germ cell populations, uniform manifold approximation and projection (UMAP) analysis identified six distinct metabolic states (States 1 – 6) (**Fig. 1a**). These metabolic states exhibited strong concordance with defined developmental stages, mapping to spermatogonial stem cells (SSCs), differentiating spermatogonia (Diff), preleptotene (PreL), and leptotene (Lep) spermatocytes (**Fig. 1b**), suggesting that transcriptionally driven metabolic reprogramming is tightly coupled to meiotic initiation. *Stra8*-deficient germ cells are arrested at the leptotene stage and fail to progress to zygotene and subsequent stages^7^.

**Figure 1.**
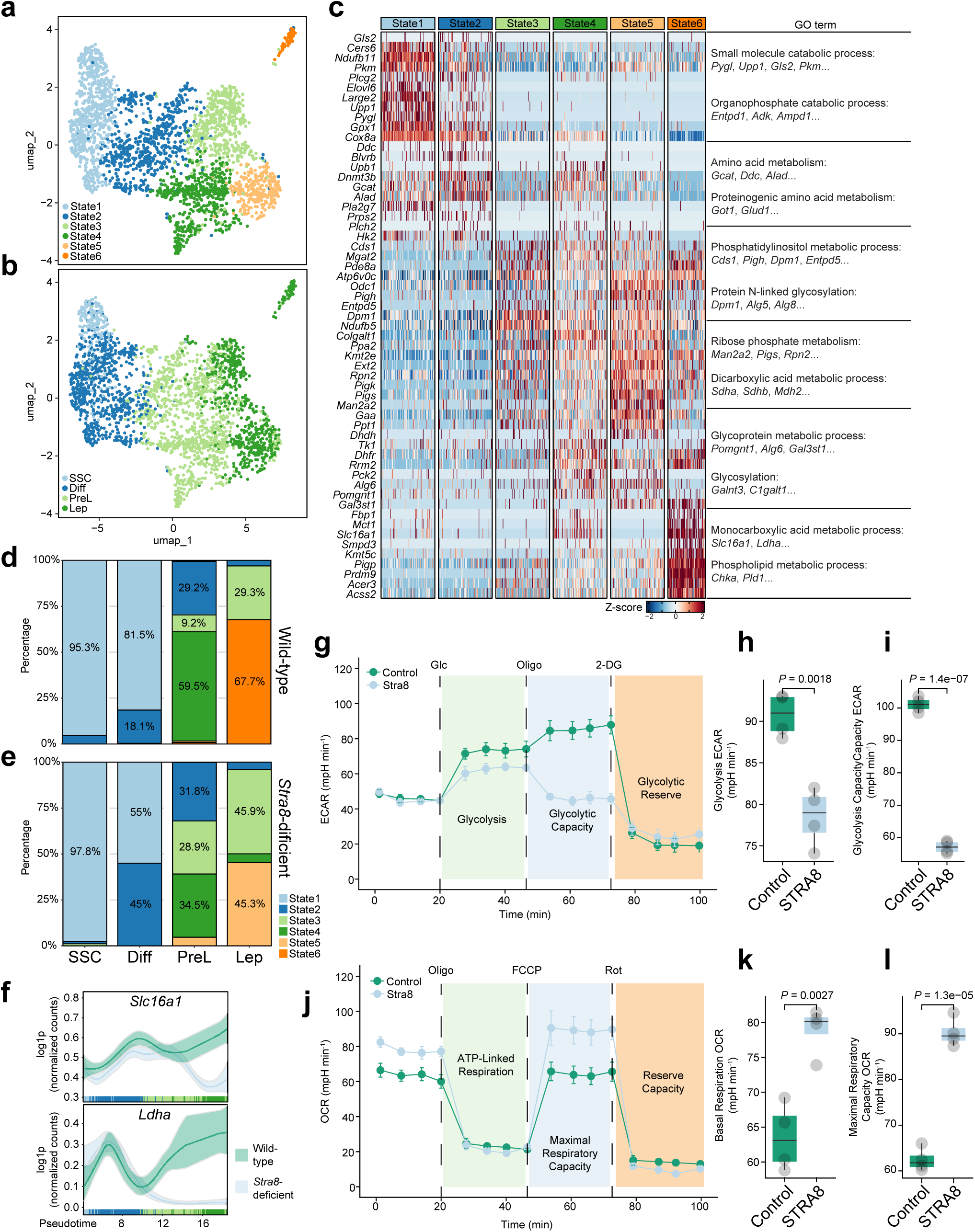
Single-cell transcriptomic analysis reveals Stra8-Dependent metabolic reprogramming in testicular cells. **a**, UMAP visualization of testicular cells clustered into six distinct states. **b**, UMAP projection colored by cell type annotation: SSC, spermatogonial stem cells; Diff, differentiating spermatogonia; PreL, preleptotene spermatocytes; Lep, leptotene spermatocytes. **c**, Heatmap of differentially expressed genes across States 1–6, with right-hand labels indicating enriched Gene Ontology (GO) terms for each gene cluster. **d**, **e**, Bar plots showing the cell state composition within each cell type in Wild-type (d) and *Stra8*-deficient (e) testes, illustrating shifts in differentiation dynamics. **f**, Pseudotime expression trajectories of *Slc16a1*and *Ldha* in testicular germ cells. **g**, Extracellular acidification rate (ECAR) profile of F9^Control^ and F9^Stra8^ cells measured by Seahorse assay. **h**, **i**, Quantification of glycolysis (h) and glycolytic capacity (i) derived from ECAR profiles, showing decreased glycolytic activity in Stra8-overexpressing cells. **j**, Oxygen consumption rate (OCR) profile of F9^Control^ and F9^Stra8^ cells. **k**, **l**, Quantification of basal respiration (k) and maximal respiratory capacity (l) derived from OCAR profiles, showing enhanced mitochondrial respiration in Stra8-expressing cells. Data are presented as mean ± s.e.m. P values were determined by two-tailed Student’s *t*-test.

To define the molecular features of these metabolic states, we performed differential gene expression analysis across states. Heatmap visualization revealed a highly stage-specific metabolic gene network (**Fig. 1c**). State 1 was enriched for small-molecule catabolic processes (*Pygl*, *Upp1*, *Gls2*, *Pkm*, etc), whereas State 2 showed enrichment for amino acid metabolism (*Gcat*, *Ddc*, *Alad*, etc). State 3 was associated with phosphatidylinositol metabolic processes (*Cds1*, *Pigh*, *Dpm1*, *Entpd5*, etc), and State 4 with ribose phosphate metabolism (*Man2a2*, *Pigs*, *Rpn2*, etc). State 5 was enriched for glycoprotein metabolic processes (*Pomgnt1*, *Alg6*, *Gal3st1,* etc), whereas State 6 was characterized by monocarboxylic acid metabolic process (*Slc16a1*, *Ldha,* etc). Interestingly, SSC, differentiating spermatogonia, and preleptotene spermatocytes in both wild-type and *Stra8*-deficent germ cells predominantly occupy State 1 – 4, consistent with the activation of amino acid, phosphatidylinositol, ribose phosphate metabolic pathways during differentiation (**Fig. 1d-e**). However, a striking divergence emerged at the onset of meiosis: wild-type cells transitioned into State 6, marked by activation of monocarboxylic acid metabolic process (**Fig. 1d**), whereas *Stra8*-deficient cells instead adopted State 5, characterized by glycoprotein metabolic processes (**Fig. 1e; Supplementary Fig. 1a-b**). Consistent with this transition, pseudotime analysis confirmed their dynamic upregulation as wild-type cells commit to the meiotic program (**Fig. 1f**). These findings identify Stra8 as a critical regulator of a metabolic switch that promotes monocarboxylic acid transport while suppressing glycoprotein metabolism during meiotic initiation.

*Slc16a1* encodes monocarboxylate transporter 1 (MCT1), a plasma membrane transporter that mediates the uptake of monocarboxylates, including lactate, across the cell membrane. *Ldha* encodes lactate dehydrogenase A (LDHA), which catalyzes the interconversion of pyruvate and lactate. Together, these factors define a core metabolic axis that regulates lactate production and utilization, suggesting that Stra8 may promote a shift toward monocarboxylic acid–dependent metabolism. To functionally validate the Stra8-associated metabolic switch, we expressed Stra8 in F9 embryonic carcinoma cells and assessed their real-time bioenergetic profiles using a Seahorse extracellular flux analyzer. Consistent with our single-cell predictions, Stra8 expression induced a profound and coordinated metabolic reprogramming. During glycolysis stress testing, Stra8-expressing cells exhibited a marked reduction in extracellular acidification rate (ECAR) (**Fig. 1g**), with both basal glycolysis and maximal glycolytic capacity significantly decreased relative to controls (**Fig. 1h-i**). In contrast, mitochondrial stress testing revealed that a robust increase in oxygen consumption rate (OCR) upon Stra8 expression (**Fig. 1j**). Stra8-expressing F9 cells showed significantly enhanced basal ATP-linked respiration and maximal respiratory capacity (**Fig. 1k, l; Supplementary Fig. 1c**). These findings demonstrate a shift away from glucose-dependent glycolysis toward oxidative metabolism, consistent with the transcriptional activation of monocarboxylic acid metabolic pathways observed in our single-cell analyses.

### *In silico* perturbation analysis links monocarboxylic acid metabolic genes to the meiotic gene regulatory network

To assess whether monocarboxylic acid metabolic genes regulate meiotic gene expression during meiotic prophase I, we employed a computational modeling approach. Specifically, we used Geneformer^19^, a context-aware deep learning model pre-trained on a large-scale single-cell transcriptomic data (**Fig. 2a**). The model was fine-tuned using our testis scRNA-seq dataset^7^ to distinguish between spermatogonial stem cells, differentiating spermatogonia, preleptotene, and leptotene spermatocytes. The fine-tuned model achieved high classification accuracy, as demonstrated by strong concordance between true and predicted cell identities in the confusion matrix and high-confidence prediction scores in out-of-sample heatmap (**Fig. 2b-c; Supplementary Fig. 2a-b**). These results establish a robust computational framework for modeling germ cell state transitions.

**Figure 2.**
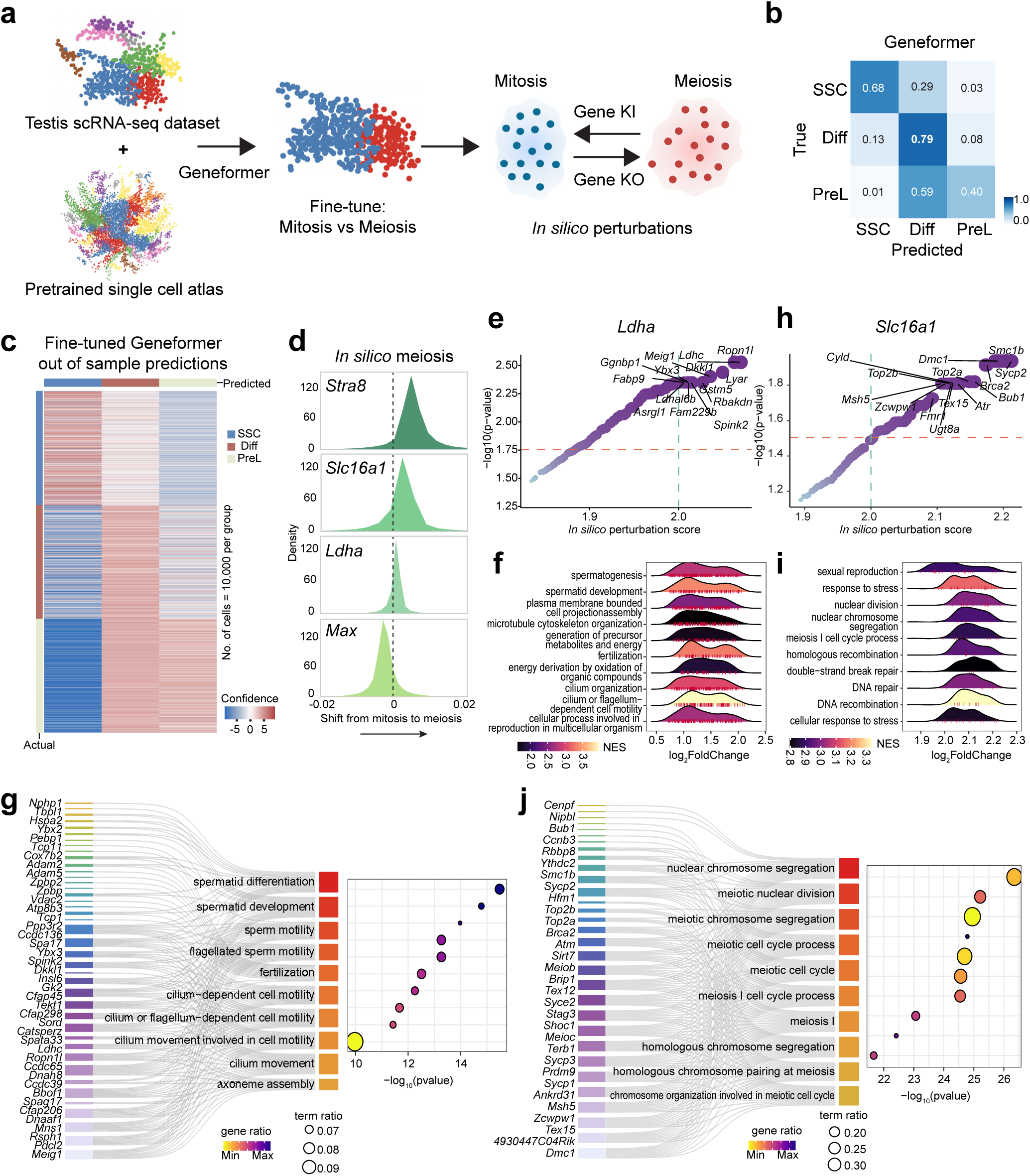
Geneformer predicts the impact of *Ldha* and *Slc16a1* perturbations on meiotic progression and related biological pathways. **a**, Schematic workflow of Geneformer fine-tuning and in silico perturbation analysis. A testis scRNA-seq dataset and a pretrained single-cell atlas are used to fine-tune. The fine-tuned model then simulates gene knock-in (KI) or knock-out (KO) to predict shifts between mitotic and meiotic states. **b**, Confusion matrix for Geneformer predictions on testicular cell types. This heatmap displays the accuracy of the fine-tuned Geneformer model in classifying SSC, Diff, and PreL cells. Rows represent true labels, and columns represent predicted labels. High diagonal values indicate good prediction accuracy. **c**, Heatmap of Geneformer’s sample predictions. The color intensity reflects the model’s prediction strength. **d**, Density plots showing in silico shifts from mitosis to meiosis upon gene perturbations. These plots illustrate the predicted shift in cell state upon in silico perturbation of *Stra8*, *Slc16a1*, *Ldha*, and *Max*. **e**, *Ldha* in silico perturbation effects on gene expression. Rank plot showing genes whose expression is significantly altered by in silico perturbation of *Ldha*, ranked by perturbation score. Genes with high perturbation scores and significance are highlighted. **f**, GSEA of pathways affected by *Ldha* in silico perturbation. Ridge plots display the Normalized Enrichment Score distribution for various biological pathways affected by *Ldha* perturbation. **g**, Enriched biological pathways and associated genes upon *Ldha* in silico perturbation. A gene-concept network plot showing the top enriched biological pathways related to spermatid differentiation, sperm motility, and cilium movement, linked to specific genes whose expression is altered by *Ldha* perturbation. **h**, *Slc16a1* in silico perturbation effects on gene expression. Rank plot showing genes whose expression is significantly altered by in silico perturbation of *Slc16a1*, ranked by perturbation score. Key meiotic genes are highlighted. **i**, GSEA of pathways affected by *Slc16a1* in silico perturbation. Ridge plots display the NES distribution for biological pathways affected by *Slc16a1* perturbation. **j**, Enriched biological pathways and associated genes upon Slc16a1in silico perturbation. A gene-concept network plot showing the top enriched biological pathways related to nuclear chromosome segregation, meiotic cell cycle, and homologous chromosome pairing, linked to specific genes whose expression is altered by *Slc16a1* perturbation.

Using this fine-tuned model, we next applied its attention-based architecture to perform *in silico* perturbations, which simulate modulation of specific metabolic factors and assessing their impact on the transition from mitosis to meiosis. As positive controls, consistent with its established role as a gatekeeper of meiotic initiation^2,5^, *in silico* activation of *Stra8* resulted in a strong shift in cell embeddings toward the meiotic state. In contrast, *in silico* activation of *Max*, which encodes an inhibitor of meiotic entry^20^, shifted cell embeddings away from the meiotic state and toward mitosis.

Notably, *in silico* activation of *Slc16a1* and *Ldha* similarly shifted cell embeddings toward the meiotic state (**Fig. 2d**). These computational predictions suggest that *Slc16a1* and *Ldha* expression contributes to the entry or progression of cells into meiosis.

To characterize the downstream transcriptional programs associated with MCT1 (encoded by *Slc16a1*) and Ldha, we analyzed gene regulatory changes induced by *in silico* perturbations of *Slc16a1 and Ldha*. *In silico* perturbation of *Ldha* identified significant enrichment of genes critical for late-stage spermatogenesis and structural maturation, including *Ggnbp1*, *Meig1*, and *Ldhc*. Gene set enrichment analysis (GSEA) further revealed that *Ldha*-perturbed transcriptomes were enriched for Gene Ontology (GO) terms associated with cilium- or flagellum-dependent cell motility, spermatid development, and axoneme assembly (**Fig. 2e-f**). Network analysis highlighted a tight linkage between *Ldha* and genes governing spermatid differentiation and development (**Fig. 2g**). These findings suggest that *Ldha*-mediated metabolic flux is primarily linked to supporting the bioenergetic and structural demands of spermiogenesis and motility acquisition, rather than early meiotic events. That’s consistent with the relatively modest shift toward the meiotic state observed upon *Ldha* activation compared with *Slc16a1* activation (**Fig. 2d**).

In contrast, *in silico* perturbation of *Slc16a1* revealed a regulatory landscape fundamentally distinct from that of *Ldha*. The top-ranked downstream targets of *Slc16a1* included canonical meiotic prophase genes, such as *Sycp3*, *Sycp1*, *Dmc1*, and *Hormad1*. Functional enrichment analysis confirmed that *Slc16a1* activity is strongly associated with meiosis I cell cycle process, homologous chromosome pairing, and double strand break repair (**Fig. 2h-i**). Furthermore, network analysis highlighted a tight linkage between *Slc16a1* and genes governing nuclear chromosome segregation and meiotic nuclear division (e.g., *Top2a*, *Smc1b*, *Rad21l1*) (**Fig. 2j**).

To further assess the regulatory role of *Slc16a1* in early meiosis, we examined the correlation between *Slc16a1* and *Ldha* expression and a curated set of 165 early meiosis-related genes^6^ using the Genotype-Tissue Expression (GTEx) database^21^ (**Supplementary Fig. 3a**). Notably, intersection analysis revealed that the top-ranked genes identified by the *in silico* perturbation analysis of *Slc16a1* significantly overlapped with these endogenously correlated genes (19 overlapping genes, including *Dmc1*, *Sycp1*, and *Prdm9*) (**Supplementary Fig. 3b**). For example, *Slc16a1* expression levels showed a robust positive correlation with both *Prdm9*, encoding a meiotic histone methyltransferase (R=0.41), and *Sycp1*, encoding a synaptonemal complex component (R=0.41) (**Supplementary Fig. 3c**). Collectively, these *in silico* genetic interaction screens suggest that *Slc16a1* contributes to the establishment of competence for meiotic entry and chromosomal recombination.

### Germline *Slc16a1* is essential for the completion of meiotic prophase

To experimentally characterize the physiological roles of MCT1 during spermatogenesis, we generated germ cell-specific *Slc16a1* conditional knockout mice by crossing *Slc16a1^flox/flox^* mice with Stra8-Cre mice (Stra8-Cre;Slc16a1^flox/flox^ mice; referred hereafter as Slc16a1^gcKO^). Stra8-Cre;Slc16a1^flox/+^ mice were used as controls.

Immunofluorescence staining confirmed that MCT1 expression was reduced in the Slc16a1^gcKO^ testis (**Supplementary Fig. 4a**). Slc16a1^gcKO^ mice appeared grossly normally, with no significant differences in body weight compared to controls. However, adult Slc16a1^gcKO^ mice consistently showed smaller testes (data not shown).

Histological examination of control and Slc16a1^gcKO^ testes revealed a complete arrest of meiosis and spermatogenesis. Control testes show normal spermatogenesis progressing to spermatid (**Fig. 3a**). In Slc16a1^gcKO^ testes, germ cells in approximately 80% of seminiferous tubules were arrested at the pachytene stage, while the remaining 20% of seminiferous tubules exhibited a complete loss of germ cells (**Fig. 3a**).

**Figure 3.**
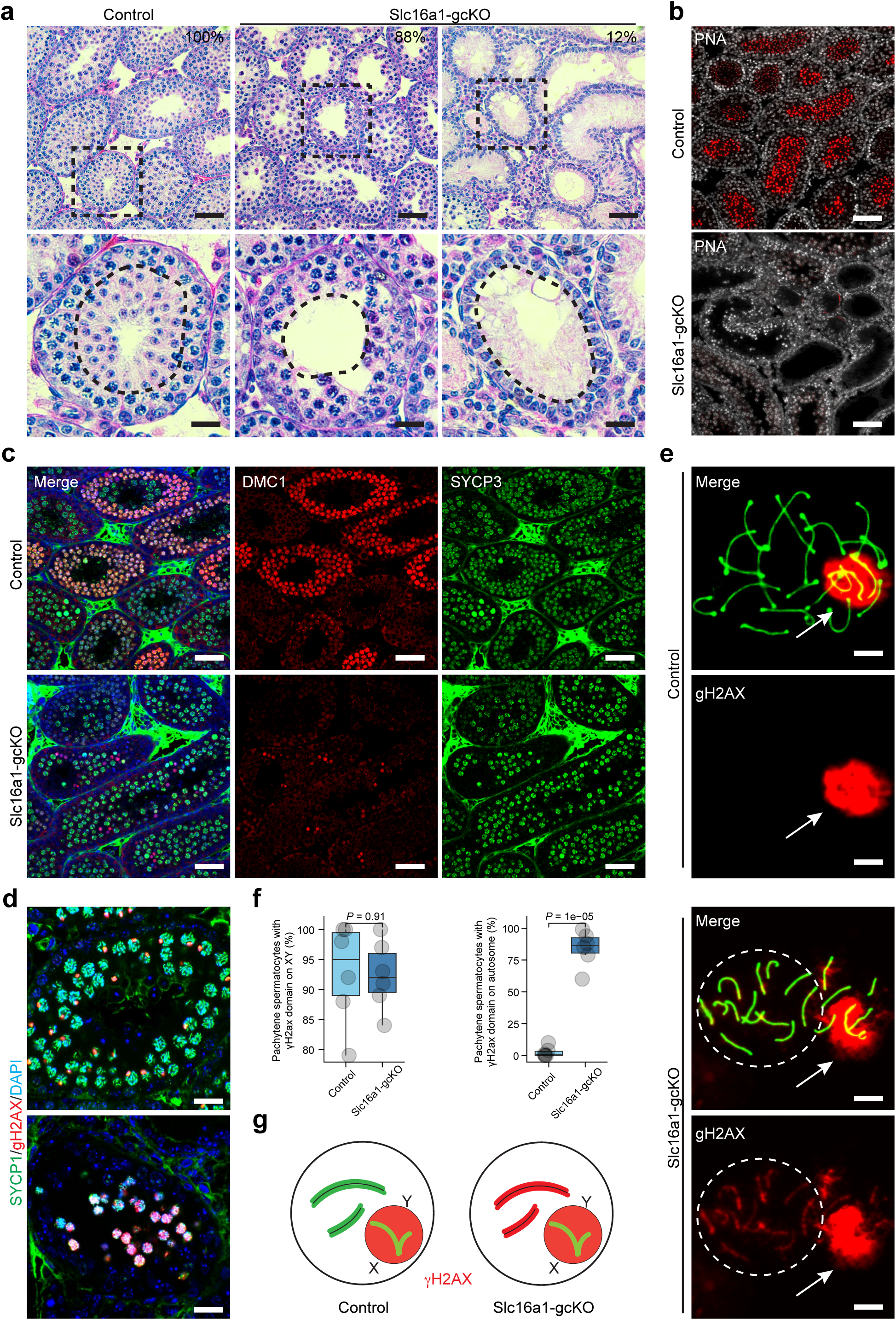
Germ cell-specific deletion of *Slc16a1* causes meiotic arrest and defective spermiogenesis. **a**, Representative H&E-stained testis sections from control and Slc16a1-gcKO mice. Percentages indicate the frequency of tubules displaying the observed phenotypes. Scale bars, 50 μm. **b**, PNA staining (red) showing impaired acrosome formation and lack of spermatid development in Slc16a1-gcKO testes compared to controls. Scale bars, 50 μm. **c**, Immunofluorescence staining of DMC1 (red) and SYCP3 (green) in control and Slc16a1-gcKO testes, highlighting defects in meiotic recombination and synapsis. Scale bars, 50 μm. **d**, Immunofluorescence staining of SYCP1 (green) and γH2AX (red) showing meiotic prophase progression. Scale bars, 20 μm. **e**, High-magnification images of pachytene spermatocytes stained for γH2AX (red) and DAPI (blue). Arrows indicate the sex body. Note the aberrant autosomal γH2AX signal in Slc16a1-gcKO spermatocytes, contrasted with the sex body-restricted signal in controls. Scale bars, 5 μm. **f**, Quantification of pachytene spermatocytes with γH2AX signal restricted to the XY body (left) or present on autosomes (right). Data are mean ± s.d. P values derived from Student’s t-test. **g**, Schematic illustrating the sex body-restricted γH2AX pattern in controls versus aberrant autosomal γH2AX localization in Slc16a1-gcKO pachytene spermatocytes, indicative of unrepaired DNA double-strand breaks.

At 8 weeks of age, control epididymides contain spermatid, whereas Slc16a1^gcKO^ epididymides show a complete absence of spermatid, confirming spermatogenic arrest (**Supplementary Fig. 4b**).

To further confirm the absence of spermatids, we performed staining with peanut agglutinin (PNA). PNA robustly labels the acrosome of round and elongating spermatids and is widely used as a marker of post-meiotic germ cell differentiation. In control testes, PNA staining revealed both punctate (dot-like) signals consistent with early proacrosomal vesicles and strong cap-like staining marking developing acrosomes in round and elongating spermatids (**Fig. 3b**). In contrast, cap-like staining was completely absent in Slc16a1^gcKO^ testes, while punctate signals were barely visible, indicating a failure to generate post-meiotic germ cells (**Fig. 3b**).

Given the meiotic arrest, we examined control and Slc16a1^gcKO^ testes at 4 weeks of age, when germ cells in meiotic prophase I are enriched. We used DMC1, a marker of meiotic recombination foci associated with programmed DNA double-strand break repair, and SYCP3, a structural component of the synaptonemal complex that marks chromosome axis formation during meiotic prophase I. We found that, in contrast to germ cells in control testes, germ cells in Slc16a1^gcKO^ testes showed a profound reduction in DMC1 signal, while SYCP3 staining appeared grossly normal (**Fig. 3c**).

DMC1-mediated recombination repair is required for progression to pachytene. To assess pachytene-stage cells, we examined γH2AX, which becomes restricted to the sex chromosomes to form the sex body during this stage. In control testes, germ cells display γH2AX localization to discrete sex bodies, consistent with pachytene progression (**Fig. 3d**). In contrast, Slc16a1^gcKO^ testes contain only germ cells with diffuse nuclear γH2AX staining, indicative of persistent DNA damage and failure to progress to pachytene (**Fig. 3d**). To confirm this result, we performed chromosome spread analysis (**Fig. 3e**). In control germ cells, γH2AX was restricted to sex bodies; however, in Slc16a1^gcKO^ germ cells, γH2AX was detected on both sex chromosomes and autosomes, suggesting unresolved DNA double-strand breaks and defective meiotic progression (**Fig. 3e–g**).

Further immunofluorescence analysis of chromosome spread analysis revealed that the early stages of homologous recombination are profoundly disrupted in Slc16a1^gcKO^ mice. In control spermatocytes, DMC1 and RAD51 foci were highly abundant along the axial elements during the zygotene stage (averaging ∼250 foci per nucleus) and were progressively resolved as cells advanced to the pachytene stage (**Fig. 4a–d**). Interestingly, this dynamic recruitment was significantly in the absence of MCT1. In Slc16a1-gcKO spermatocytes, the numbers of DMC1 and RAD51 foci were drastically reduced at the zygotene stage and remained low during the pachytene stage (**Fig. 4a–d**). Due to the successful processing of DSBs by RAD51 and DMC1 is an absolute prerequisite for the maturation of recombination intermediates into crossovers, we assessed the downstream formation of chiasmata. We quantified the localization of MLH1, a mismatch repair protein that strictly marks prospective crossover sites in late pachytene cells. Consistent with the upstream failure in homologous recombination initiation, the number of MLH1 foci per nucleus was significantly diminished in Slc16a1-gcKO pachytene spermatocytes compared to the robust 22–25 foci typically observed in controls (**Fig. 4e–f**). These data provide the direct cytological basis for the meiotic failure observed in Slc16a1-gcKO germ cells.

**Figure 4.**
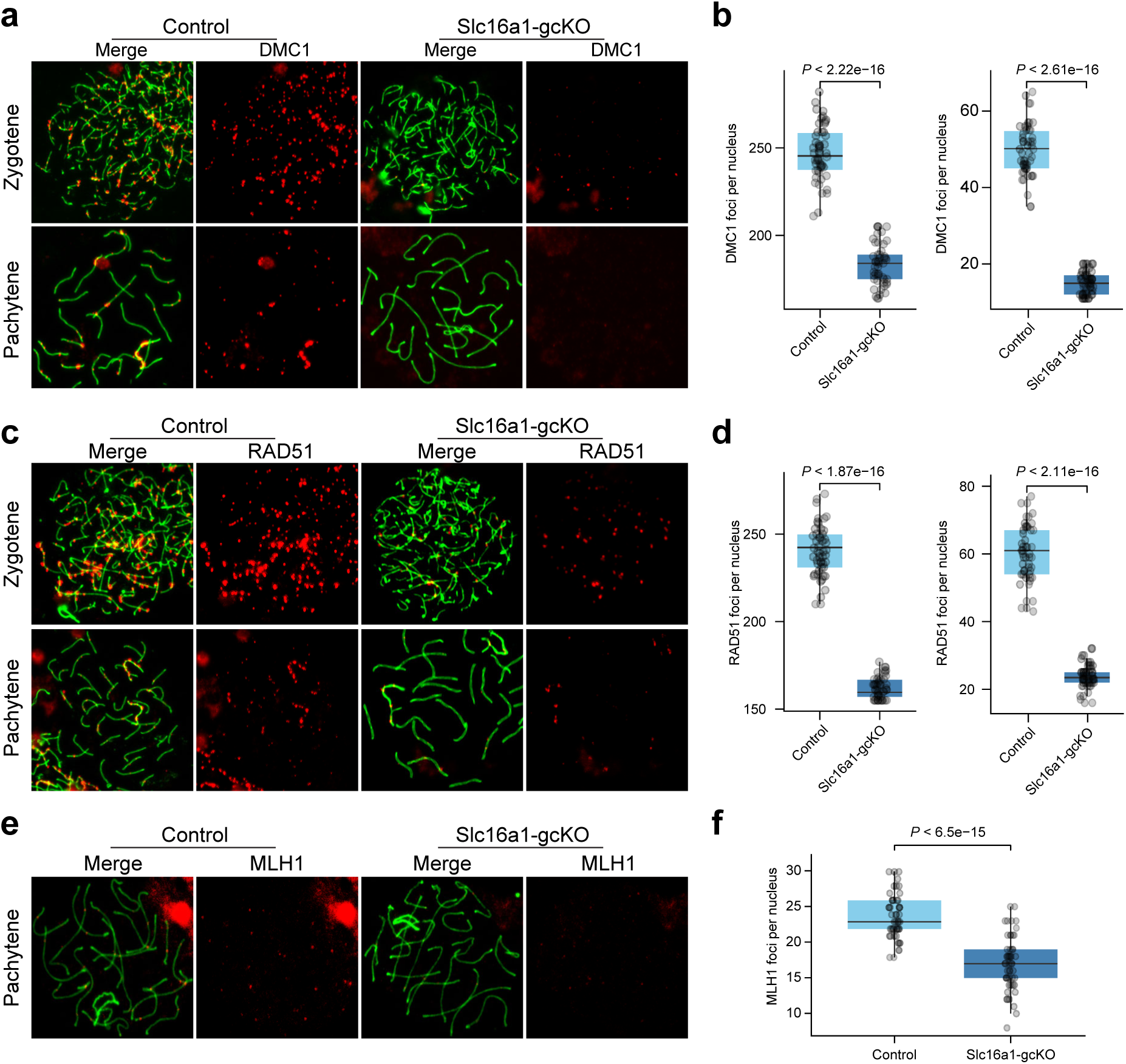
*Slc16a1* deficiency impairs homologous recombination and crossover formation during meiosis. **a**, Immunofluorescence images of DMC1 (red) and SYCP3 (green) in zygotene and pachytene spermatocytes from control and Slc16a1-gcKO mice. **b**, Box plots show a significant reduction in DMC1 foci per nucleus in Slc16a1-gcKO spermatocytes at both stages. **c**, Immunofluorescence staining of RAD51 (red) foci on SYCP3-stained (green) chromosome axes. **d**, Box plots show a significant reduction in RAD51 foci per nucleus in Slc16a1-gcKO spermatocytes at both stages. **e**, Immunofluorescence staining of MLH1 (red) foci in pachytene spermatocytes, marking designated crossover sites. **f**. Slc16a1-gcKO cells display a significant reduction in MLH1 foci compared to controls. All quantitative data are presented as box plots. P values were determined by two-tailed Student’s *t*-test. SYCP3 marks the synaptonemal complex.

Inside the seminiferous tubules, because *Slc16a1* expression is detected in Sertoli cells (**Supplementary Fig. 5a**), we generated a Sertoli cell–specific *Slc16a1* knockout using Amh-Cre mice (Slc16a1^scKO^). In contrast to the Slc16a1-gcKO, Slc16a1-scKO mice did not show impaired meiosis (**Supplementary Fig. 5b**). However, the seminiferous tubules of Slc16a1-scKO mice lack post-meiotic spermatids (**Supplementary Fig. 5b**). Consistent with this finding, examination of the cauda epididymis in Slc16a1-scKO mice revealed significant luminal defects: unlike controls, which contained abundant healthy spermatozoa, the epididymal tubules of Slc16a1-scKO mice were filled with aberrant cellular debris and abnormal sperm aggregates (**Supplementary Fig. 5c**). These data suggest that MCT1 expression in Sertoli cells is critical for downstream sperm maturation.

### *Slc16a1* is required to sustain the meiotic transcriptional program

To elucidate the mechanism underlying the meiotic arrest observed in Slc16a1-gcKO testes, we performed RNA-sequencing (RNA-seq) from control and Slc16a1-gcKO testes at postnatal day 18 (n = 2 per group). At this stage, germ cell composition is relatively stable between control and Slc16a1-gcKO testes, despite arrest in pachytene. Principal component analysis (PCA) revealed that the transcriptomes of Slc16a1-gcKO germ cells were distinct from those of control, despite some variability among control biological replicates (**Fig. 5a**). Volcano plot analysis identified a large number of differentially expressed genes (DEGs), with a predominant fraction significantly downregulated in Slc16a1-gcKO testes, suggesting a failure to activate or maintain the specific transcriptional program required for meiosis (**Fig. 5b**). We further characterized the identity of the downregulated genes by crossing them with stage-specific expression profiles from published dataset of normal spermatogenesis^22,23^. Interestingly, the genes downregulated in Slc16a1-gcKO testis are typically those with peak expression during the pachytene spermatocytes (PS) and round spermatid (RS) stages (**Fig. 5c**).

**Figure 5.**
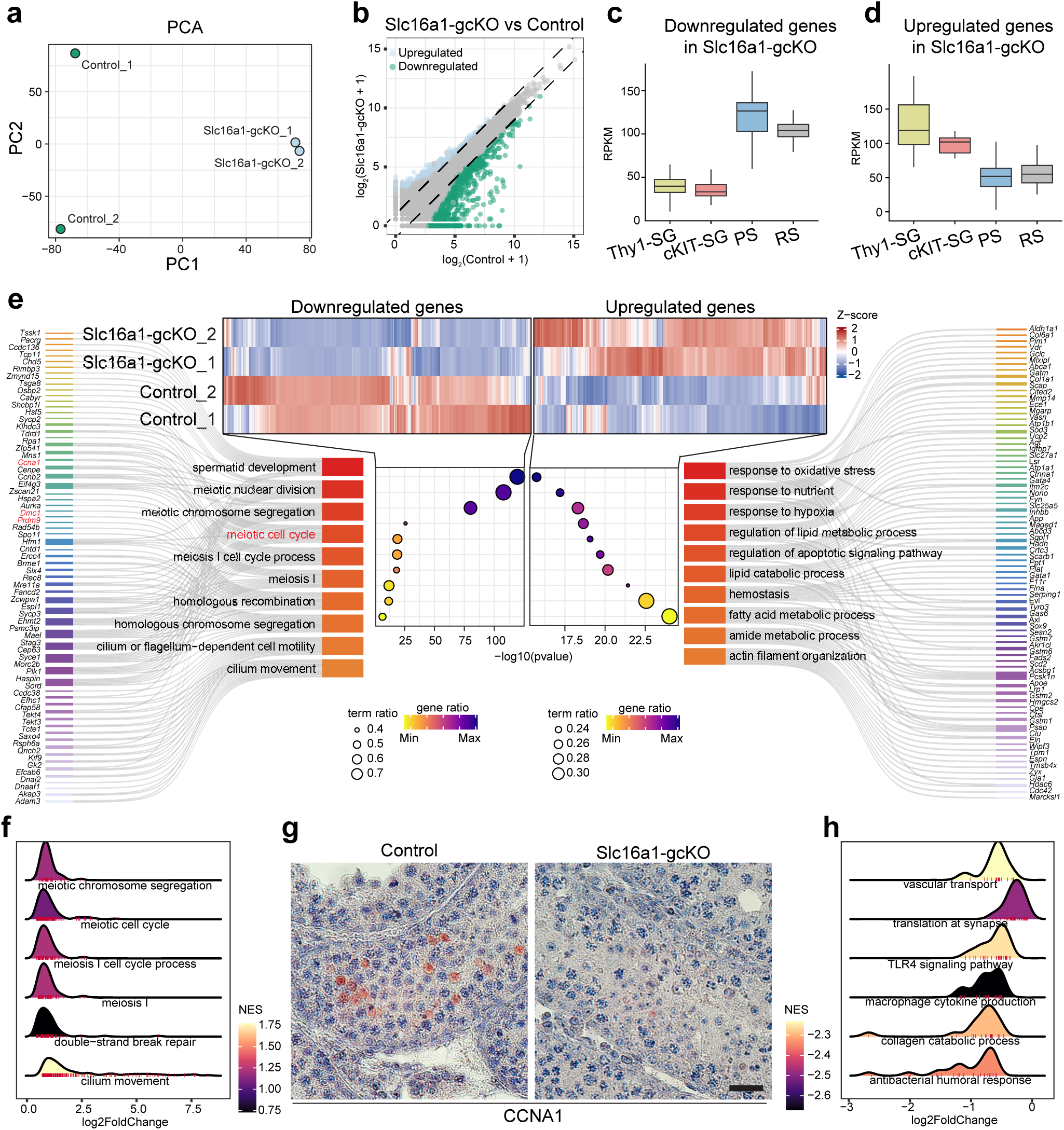
Transcriptomic Analysis of Slc16a1-gcKO Testes Reveals Disrupted Meiotic Defect by Meiotic Cell Cycle Arrest through CCNA1. **a**, PCA analysis of RNA-seq data from control and Slc16a1-gcKO testes showed clear separation between control and Slc16a1-gcKO samples. **b**, Scatter plot of gene expression comparing Slc16a1-gcKO vs. control testes displayed the differentially expression level of all genes. **c**, Box plot of RPKM values for representative downregulated genes in published dataset. **d**, Box plot of RPKM values for representative upregulated genes in published dataset. **e**, Heatmap of differentially expressed genes in Slc16a1-gcKO testes. This heatmap displayed the z-score normalized expression of all genes that are either upregulated in control testes or upregulated in Slc16a1-gcKO testes. GO analysis pathways downregulated in Slc16a1-gcKO testes. And GO analysis pathways upregulated in Slc16a1-gcKO testes. **f**, GSEA of downregulated pathways in Slc16a1-gcKO testes. Ridge plots show the NES distribution for key biological pathways that are significantly downregulated in Slc16a1-gcKO testes. **g**, Immunohistochemistry staining for CCNA1 (Cyclin A1) in control and Slc16a1-gcKO testes showed CCNA1 protein expression (brown) is visibly reduced in Slc16a1-gcKO testes. **h**, GSEA of upregulated pathways in Slc16a1-gcKO testes. Ridge plots show the NES distribution for key biological pathways that are significantly upregulated in Slc16a1-gcKO testes.

Functional enrichment analysis revealed that these genes are associated with meiotic cell cycle, homologous recombination and meiotic chromosome segregation (**Fig. 5d**). Specifically, genes encoding core meiotic regulators, including *Dmc1*, *Sycp2*, *Rec8* and *Hormad1*, showed a profound reduction in gene expression (**Fig. 5d**). This transcriptomic signature is consistent with our observation that Slc16a1-gcKO germ cells arrest at a pachytene-like stage and fail to initiate the post-meiotic program (**Fig. 4**). GSEA analysis shows enrichment for meiosis-related pathways, e.g., meiotic chromosome segregation, meiotic cell cycle, meiosis I cell cycle process (**Fig. 5f**)

In contrast, genes upregulated in Slc16a1-gcKO germ cells are enriched for markers associated with undifferentiated spermatogonia (Thy1-SG) (**Fig. 5d**). GO analysis revealed enrichment for pathways associated with oxidative stress, response to nutrient, and response to hypoxia (**Fig. 5e**). We also observed the activation of pathways associated with lipid metabolic processes and apoptotic signaling, suggesting that impaired lactate transport due to *Slc16a1* deletion imposes metabolic stress that triggers a starvation-like response and ultimately leads to germ cells loss. GSEA analysis shows enrichment for functions such as vascular transport, translation at synapse, TKR4 signaling pathway (**Fig. 5h**)

Among the DEGs between control and Slc16a1-gcKO testes, *Ccna1*, which encodes Cyclin A1 (CCNA1), was notably downregulated in Slc16a1-gcKO testes (**Fig. 5h**). CCNA1 is an essential, meiosis-specific cyclin required for the transition from pachytene to diplotene stage, and its genetic ablation results in that phenocopies the meiotic arrest observed in Slc16a1-gcKO mice^24^. To validate the RNA-seq results, immunohistochemical staining of CCNA1 showed robust CCNA1 signal in the expected cell layers (pachytene) of control seminiferous tubules where cells accumulate, whereas CCNA1 expression was virtually absent in Slc16a1-gcKO tubules. This molecular evidence, together with the widespread downregulation of key meiotic genes, indicate that MCT1 is required for germ cells to progress through the pachytene stage.

### MCT1 epigenetically licenses meiotic transcription via histone lactylation

MCT1 functions primarily as an importer of monocarboxylic acids, including lactate; however, accurately measuring lactate levels in the testis is challenging due to its heterogenous cellular composition, which includes both somatic and germ cells spanning multiple mitotic and meiotic stages. We therefore used histone lactylation, a lactate-induced epigenetic modification^17^, as a proxy for intracellular lactate availability. Recently, we reported that histone lactylation correlates with meiotic gene expression during meiotic prophase^18^. We hypothesized that loss of MCT1 would impair histone lactylation and consequently reduce meiotic gene expression.

We examined histone lysine (K) lactylation at H3K9, H3K14, H3K18, H4K5, H4K8, H4K12, and H4K16. Immunohistochemical staining revealed a robust and specific accumulation of H3K14la, H3K18la, K4K8la, H4K12, and H4K16la in the nuclei of wild-type pachytene spermatocytes (**Fig. 6a, b; Supplementary Fig. 6**). Interestingly, H3K14la, H4K18la, H4K12la, and H4K16la signals were markedly reduced in the Slc16a1-gcKO seminiferous tubules. These data suggest that MCT1-imported lactate is a substrate driving histone lactylation during meiotic prophase (**Fig. 6a, b**).

**Figure 6.**
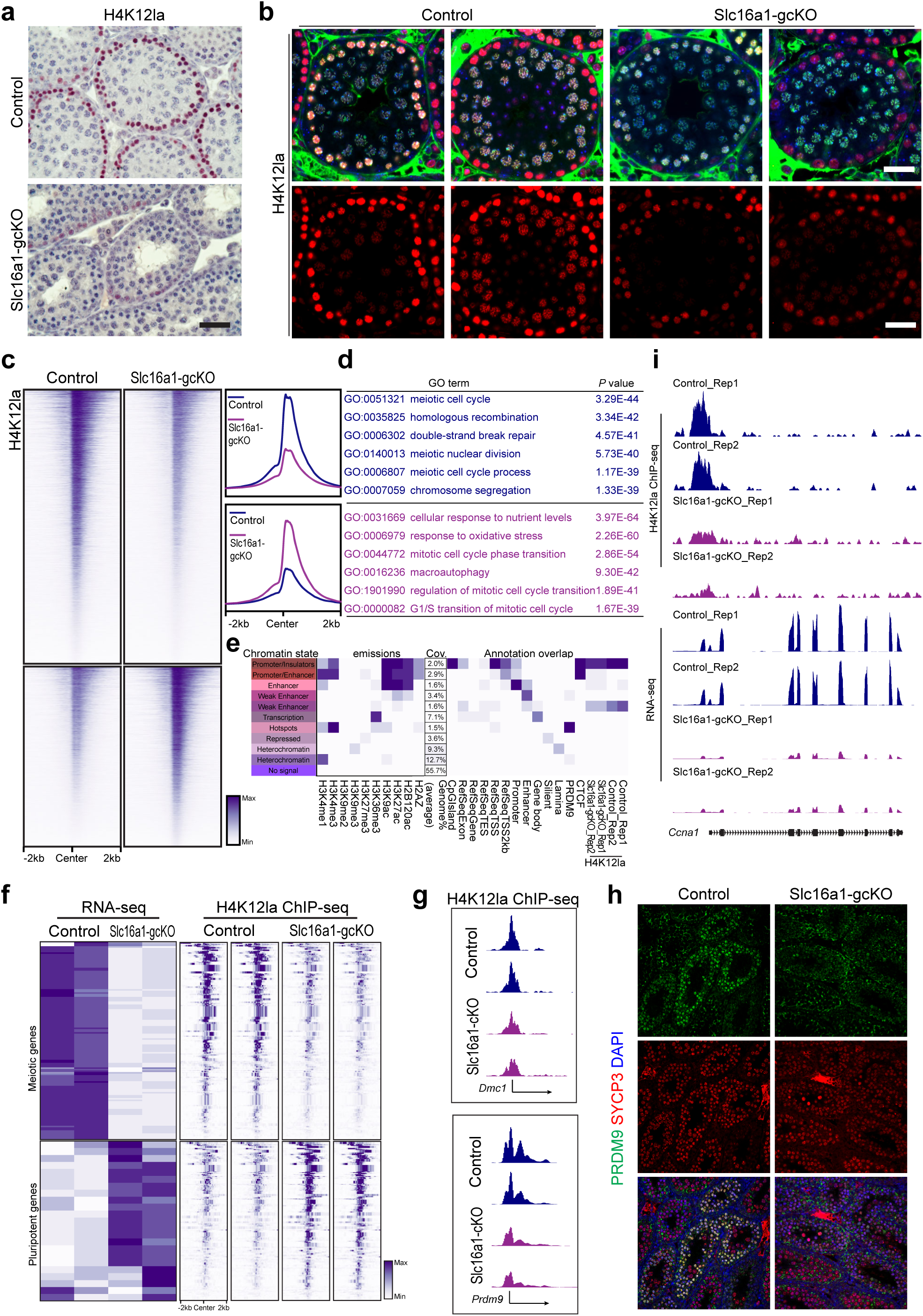
Slc16a1-dependent H4K12 lactylation regulates meiotic gene expression. **a**, Immunohistochemistry staining of H4K12la in control and Slc16a1-gcKO mouse testes, showing reduced H4K12la levels in the Slc16a1-gcKO group. **b**, Immunofluorescence staining of H4K12la (red) in control and Slc16a1-gcKO testes. Scale bars, 20 μm. **c**, Heatmaps and average profiles of H4K12la ChIP-seq signal in control and Slc16a1-gcKO testes, revealing a global reduction of H4K12la enrichment. **d**, Gene Ontology (GO) enrichment analysis of genomic regions with differential H4K12la occupancy. **e**, Chromatin state annotation integrating histone modifications and H4K12la occupancy. **f**, Heatmap correlating RNA-seq expression levels and H4K12la ChIP-seq signals for meiotic and pluripotent gene sets. **g**, **h**, Genome browser tracks of H4K12la ChIP-seq at *Dmc1* and *Prdm9* loci (left), and immunofluorescence staining of PRDM9 (green) and SYCP3 (red) in control and Slc16a1-gcKO testes (right). **i**, Genome browser tracks of H4K12la ChIP-seq and RNA-seq at the *Ccna1* locus, showing decreased H4K12la binding and mRNA expression in Slc16a1-gcKO testes.

To map the genome-wide H4K12la landscape and understand its regulatory scope, we performed chromatin immunoprecipitation followed by sequencing (ChIP-seq). In wild-type testes, H4K12la signals were highly enriched at the transcription start sites (TSS) and active promoter/enhancer regions across the genome (**Fig. 6c, e**).

Gene Ontology (GO) analysis of the H4K12la-bound loci in wild-type cells showed enrichment for biological processes essential for meiosis, including meiotic cell cycle, homologous recombination and double-strand break repair (**Fig. 6d**). However, in Slc16a1-gcKO testes, the H4K12la peaks at these meiotic loci were profoundly reduced. Instead, the residual H4K12la signals aberrantly shifted towards stress-related pathways, such as cellular response to nutrient levels and response to oxidative stress (**Fig. 6d**), which mirrors the starvation and stress response signatures observed in our transcriptomic analysis (**Fig. 5e**).

By integrating our H4K12la ChIP-seq and RNA-seq datasets, we uncovered a direct epigenetic mechanism governing meiotic progression. Global heatmap profiling confirmed that the broad panel of meiotic genes strongly depends on H4K12la deposition for their active expression, a signature that is completely lost upon *Slc16a1* deletion (**Fig. 6f**). Specific examination of key meiotic drivers and homologous recombination factors, including *Dmc1* and the meiotic histone methyltransferase *Prdm9*, revealed prominent H4K12la peaks at their promoter regions in control samples, which were lost in Slc16a1-gcKO samples (**Fig. 6g**). Consistently, both DMC1 and Prdm9 expression is profoundly lost in the Slc16a1-gcKO testis (**Fig. 3c**; **Fig. 6h**).

Additionally, *Ccna1*, which encodes the essential late-prophase cell-cycle regulator that we identified as being profoundly downregulated in the mutant (**Fig. 5h**), displayed a robust H4K12la peak at its promoter in control samples that was markedly reduced in Slc16a1-gcKO samples (**Fig. 6i**).

## DISCUSSION

The transition from mitosis to meiosis requires coordinated regulation of transcriptional, chromosomal, and metabolic programs^25,26^. While transcriptional regulators such as STRA8 and MEIOSIN have been well defined^1,4,5^, how metabolic inputs are integrated into meiotic progression has remained unclear. Here, we identify a previously unrecognized, germ cell–intrinsic MCT1-dependent metabolic checkpoint at the pachytene stage. Germ cell–specific deletion of *Slc16a1* results in a stringent pachytene arrest, accompanied by widespread transcriptional and epigenetic dysregulation. These findings establish MCT1-dependent monocarboxylate transport as an instructive metabolic input that epigenetically licenses the transcriptional programs required for meiotic progression.

Our data show that germ cell–specific deletion of *Slc16a1* results in defective homologous recombination, persistent DNA damage, and failure to activate key meiotic regulators, including *Dmc1*, *Prdm1*, and *Ccna1*. These phenotypes are hallmarks of pachytene checkpoint failure, a surveillance mechanism that ensures proper recombination and chromosomal integrity before progression^9^. Importantly, our findings extend the concept of the pachytene checkpoint beyond chromosomal events by identifying metabolic state as a critical determinant of checkpoint passage. Rather than being solely a consequence of failed recombination, the arrest observed in *Slc16a1*-deficient germ cells suggests that insufficient monocarboxylate transport imposes a metabolic constraint that prevents completion of the meiotic program. Thus, progression through pachytene appears to require not only successful chromosomal events but also an appropriate metabolic state. We further link this metabolic state to pachytene regulation through histone lactylation–mediated epigenetic control of meiotic gene expression. However, we cannot exclude possibility that the observed arrest is secondary to upstream meiotic defects caused by loss of MCT1. Distinguishing between a direct metabolic checkpoint and an indirect consequence fo recombination failure will require further investigation.

Our integrative analyses reveal that STRA8 drives a metabolic switch during meiotic initiation, promoting a transcriptional program enriched for monocarboxylic acid metabolism. In contrast, *Stra8*-deficient germ cells fail to activate this program and instead adopt an alternative metabolic state that predominantly features glycoprotein metabolic process, consistent with their inability to progress beyond early meiotic stages. These findings suggest that metabolic reprogramming is not merely downstream of meiotic entry but is an integral component of the fate transition itself. In this framework, MCT1 represents a key effector linking STRA8-dependent transcriptional programs to metabolic flux required for meiotic progression.

A major mechanistic insight from this study is that MCT1-dependent transport is required to sustain histone lactylation, particularly H4K12la, at promoters of meiotic genes. Loss of MCT1 in germ cells leads to a marked depletion of H4K12la at these loci, accompanied by redistribution of this modification toward stress-response pathways. This shift closely mirrors the transcriptional reprogramming observed in *Slc16a1*-deficient germ cells, which exhibit suppression of meiotic genes and activation of starvation- and stress-related pathways.

Accurate measurement of intracellular metabolites requires rapid snap-freezing to preserve metabolic states. However, the testis contains a highly heterogeneous mixture of somatic and germ cells at distinct developmental stages, making it technically challenging to isolate a homogeneous population without cell dissociation, followed by cell sorting, which could alter metabolic states. Thus, direct measurement of intracellular metabolites in specific germ cell populations remains technically challenging. we use histone lactylation as a functional readout of lactate as a monocarboxylate-dependent metabolic input. As lactylation is driven by lactate^17^, our data support a model in which MCT1-mediated lactate import into germ cells provides a critical metabolic substrate that enables deposition of histone lactylation marks at meiotic gene loci, thereby licensing their expression.

Our analyses further reveal that components of the monocarboxylate metabolic pathway have distinct stage-specific roles. While both *Slc16a1* and *Ldha* are associated with monocarboxylic acid metabolism, *in silico* perturbation analysis indicates that *Slc16a1* is preferentially linked to early meiotic processes, including recombination and chromosome segregation, whereas *Ldha* is more strongly associated with later stages such as spermiogenesis. This distinction highlights that metabolic pathways are functionally compartmentalized across developmental stages. In particular, MCT1-dependent transport appears uniquely required to support the transcriptional and epigenetic programs of meiotic prophase.

Our genetic studies demonstrate that the requirement for MCT1 in meiotic progression is germ cell–autonomous. Although MCT1 is expressed in both germ cells and Sertoli cells, only germ cell–specific deletion results in meiotic arrest, whereas Sertoli cell–specific deletion does not impair meiotic progression but instead results in spermaid defects. This finding indicates that the critical function of MCT1 lies within germ cells themselves, consistent with a model in which intracellular metabolic state directly regulates chromatin and transcriptional programs. While Sertoli cells likely contribute to the metabolic environment, the ability of germ cells to utilize monocarboxylate transport appears to be the limiting factor for progression through pachytene.

Several limitations should be noted. First, although histone lactylation provides a useful proxy for monocarboxylate-dependent metabolic (lactate) input, direct measurement of intracellular metabolite levels in germ cells enriched at specific stages will be important to refine this model. Second, the enzymatic machinery—specifically, histone lactyl-transferase and delactylase—responsible for adding and removing histone lactylation in germ cells remains incompletely understood and warrants further investigation. Our past study has identified HBO1 mediating H4K8la^18^. It is unclear whether HBO1 is also responsible for H4K12la. Third, while our data establish a strong correlation between H4K12la and meiotic gene expression, the molecular mechanisms by which this modification promotes transcription remain to be defined. Identifying proteins that recognize or interpret histone lactylation marks will be critical for understanding how metabolic signals are translated into gene regulatory outputs.

In summary, we identify MCT1-dependent monocarboxylate transport as a critical regulator of meiotic progression and define a metabolic checkpoint at the pachytene stage. By linking metabolic flux to epigenetic regulation via histone lactylation, our findings establish a mechanistic framework through which metabolic state governs meiotic cell–fate decisions. These results broaden the conceptual framework of meiosis by positioning metabolism as an instructive determinant of germ cell development.

## Materials and Methods

### Mice

All animal experiments were approved by the Institutional Animal Care and Use Committee (IACUC) at the University of Kansas Medical Center in strict accordance with its regulatory and ethical guidelines and performed in accordance with relevant guidelines. Mice were maintained on a C57BL/6 background under a 12-hour light/dark cycle with ad libitum access to food and water. Stra8-Cre mice were crossed with Slc16a1^fl/fl^ mice to generate germline-specific conditional knockout lines (Stra8-cre; Slc16a1^fl/fl^). To generate Sertoli cell-specific knockouts, Amh-Cre mice were crossed with Slc16a1^fl/fl^ mice. Genotyping was performed using PCR on tail genomic DNA with specific primers as previously described.

### Single-cell RNA-sequencing Data Analysis

Raw sequencing output was processed using the Cell Ranger pipeline (10x Genomics) with default parameters. Read alignment was performed against the mouse reference index (mm10). To facilitate comparative analysis, raw count matrices from wild-type and Stra8-deficient samples were merged, and cells were assigned unique, sample-specific identifiers to prevent duplication of non-unique barcodes across runs. Transcripts mapped to the same genes with identical unique molecular identifiers (UMIs) were collapsed. The processed raw count matrices served as the input for downstream quality control and analysis using the Seurat pipeline. Normalization, integration, and cell type assignment Seurat V4 implemented in R was utilized for cell type identification and metabolic profiling. For each sample, UMI count data were normalized using regularized negative binomial regression via the SCTransform function. Major germ cell populations in the testis were identified based on the expression of well-established markers for each developmental timepoint, including spermatogonial stem cells (SSCs), differentiating spermatogonia, and preleptotene/leptotene spermatocytes. To ensure data quality, cells exhibiting aberrant mitochondrial gene expression (indicative of stress/apoptosis) or low feature counts were filtered out. Cells displaying hybrid expression profiles characteristic of doublets were also identified and removed. This filtering strategy yielded a high-quality single-cell dataset utilized to define the metabolic state transitions during meiotic initiation. To systematically characterize the metabolic landscape during meiotic initiation, genes associated with metabolic pathways were retrieved from the Molecular Signatures Database (MSigDB) for dimension reduction and clustering analysis. Based on the expression of established germ cell markers (Plzf for SSCs, Kit/Stra8 for differentiating spermatogonia, and Sycp3/Dmc1 for meiotic spermatocytes), germ cell populations (SSCs, differentiating spermatogonia, preleptotene, and leptotene spermatocytes) were extracted from the merged wild-type and Stra8-deficient datasets. Principal component analysis (PCA) was performed using exclusively this metabolic gene set to define the cellular metabolic states. The top principal components were utilized for dimension reduction via Uniform Manifold Approximation and Projection (UMAP) and clustering based on the shared nearest neighbor (SNN) graph method in Seurat. This analysis identified six distinct metabolic states associated with specific stages of meiotic entry. Differentially expressed metabolic genes defining each state were identified using the Seurat function FindAllMarkers. To reveal the dynamic patterns of metabolic pathway activity across these states, single-sample Gene Set Enrichment Analysis (ssGSEA) was performed.

### Pseudo-time analysis

Pseudo-time trajectory analysis was performed for the temporal expression patterns of metabolic genes during the transition from mitosis to meiosis. Germ cells spanning the developmental continuum from SSCs to leptotene spermatocytes were extracted, and PCA was performed to capture the major variance associated with differentiation. The pseudo-time axis was reconstructed based on the leading principal components using the Slingshot package in R. Germ cells from wild-type and Stra8-deficient testes were ordered sequentially along this inferred trajectory. Subsequently, the dynamic expression patterns of key metabolic regulators along the pseudo-time axis were fitted and visualized using the SCP package to illustrate the metabolic switch toward anaerobic glycolysis during meiotic progression.

### Metabolic Flux Analysis

To infer metabolic activity at single-cell resolution, we performed scFEA analysis. A genome-scale metabolic model was mapped to the single-cell gene expression data to estimate the flux rates of metabolic reactions. The activity of the glycolysis-TCA cycle subnetwork was quantified, and differential flux analysis was conducted to compare the conversion rates of pyruvate to lactate (reaction M_6) between wild-type and Stra8-deficient germ cells. Flux differences with a Cohen’s D > 1.5 were considered significant. **In Silico Perturbation**

The model was fine-tuned on our testis scRNA-seq dataset to classify mitotic spermatogonia and pre-meiotic spermatocytes. To simulate gene perturbation, target genes (e.g., Mct1, Ldha, Stra8) were computationally removed from or moved to the front of the rank-value encoding of each cell’s transcriptome. The impact was quantified by calculating the cosine similarity between the original and perturbed cell embeddings. Genes shifting cell embeddings toward the meiotic state were identified, and Gene Ontology (GO) enrichment analysis was performed on the predicted downstream targets to elucidate affected biological pathways.

### Seahorse Assay

Cellular bioenergetics were measured using an XF96 Extracellular Flux Analyzer (Agilent). F9 cells stable transfected with Stra8 overexpression or control vectors were seeded onto XF96 microplates. For Glycolysis Stress Test, Extracellular Acidification Rate (ECAR) was measured following sequential injections of glucose (10 mM), oligomycin (1 μM), and 2-deoxy-glucose (50 mM). For Mito Stress Test, Oxygen Consumption Rate (OCR) was measured following injections of oligomycin (1 μM), FCCP (1 μM), and Rotenone/Antimycin A (0.5 μM). Data were normalized to cell number.

### Histology and Immunofluorescence

Testes were fixed in 4% paraformaldehyde (PFA) overnight, dehydrated, and embedded in paraffin. Sections (6 μm) were deparaffinized and rehydrated. For H&E Staining, sections were stained with hematoxylin and eosin for morphological analysis. Immunofluorescence: Antigen retrieval was performed in citrate buffer (pH 6.0). Sections were blocked with 5% BSA and incubated overnight at 4°C with primary antibodies against MCT1 (1:200), SYCP3 (1:500), SYCP1 (1:500), γH2AX (1:500), PLZF, WT1, and CCNA1. After washing, sections were incubated with Alexa Fluor-conjugated secondary antibodies. Nuclei were counterstained with DAPI. Images were acquired using a confocal microscope.

### Chromosome spread

The testes were first decapsulated by removing the tunica albuginea and subsequently transferred into 15 mL tubes containing 5 mL of TIM buffer (104 mM NaCl, 45 mM KCl, 1.2 mM MgSO₄, 0.6 mM KH₂PO₄, 6.0 mM sodium lactate, 1.0 mM sodium pyruvate, and 0.1% glucose) supplemented with 1 mg/mL collagenase IV. The samples were incubated with gentle shaking for 1 h at room temperature. Following incubation, the tissue was pelleted by centrifugation at 300 × g for 5 min at room temperature, and the supernatant was carefully removed. The tissue was then further dissociated by repeated pipetting with a 1 mL tip for 5 min to obtain a single-cell suspension, which was resuspended in 500 µL TIM buffer. Subsequently, 75 mM sucrose solution was added, and 200 µL of the cell suspension was spread onto Superfrost Plus glass slides (Fisher Scientific) pre-coated with a thin layer of 1% paraformaldehyde (PFA). The slides were allowed to partially dry in a closed slide box for 1 h, followed by further drying with the lid partially open for an additional 2 h at room temperature. Finally, the slides were washed with 0.4% Photo-Flo (Nikon), air-dried, and stored at −80 °C until use.

### Bulk RNA-sequencing

Total RNA was extracted from purified germ cells or whole testis tissue of Mct1 cKO and control mice using TRIzol reagent. RNA integrity was assessed using an Agilent Bioanalyzer (RIN > 8). Libraries were constructed using the TruSeq Stranded mRNA Library Prep Kit and sequenced on an Illumina platform (PE150). Reads were aligned to the mouse genome using STAR, and differential expression analysis was performed using DESeq2. GO enrichment analysis was conducted using the clusterProfiler R package.

### Bulk ChIP-sequencing

Chromatin immunoprecipitation (ChIP) was performed on testicular cells following removal of the tunica albuginea. Dissociated cells were crosslinked in 1% formaldehyde for 10 minutes and subsequently quenched. The tissue was washed twice with PBS prior to cell lysis in 1 ml EZ-ChIP lysis buffer (Millipore, 17-371RF). Chromatin was fragmented to an average size of ∼1000 bp by sonication using a Diagenode Pico Bioruptor. The lysate was pre-cleared with Protein G beads, and 10 µl of the resulting chromatin was reserved as input. The remaining chromatin was incubated overnight at 4°C with either H4K8la antibody or preimmune IgG as a control. Immune complexes were captured by incubation with Protein A/G agarose beads for 2 hours. Beads were washed sequentially, and chromatin was eluted in buffer containing 1% SDS and 0.1 M NaHCO₃ (pH 9.0) at 65°C. Crosslinks were reversed overnight at 65°C, followed by deproteinization at 45°C for 2 hours. DNA was then purified for downstream analysis. Enrichment was assessed by high-throughput sequencing.

ChIP-seq data processing began with adapter trimming using Cutadapt (v2.10), retaining reads with a minimum length of 25 bp. Filtered paired-end reads were aligned to the mouse reference genome (mm10) using Bowtie2 (v2.4.2) with the “--very-sensitive-local” setting. Duplicate reads were marked using Picard (MarkDuplicates), and BAM files were further filtered with SAMtools to remove unmapped reads, non-primary alignments, low-quality reads, and duplicates (-F 1804 -f 2). Peak calling was performed using MACS2 with a q-value cutoff of 0.05. Consensus peak sets for biological replicates were defined by intersecting peaks using BEDTools, retaining only peaks present in both replicates. Signal visualization was performed using deepTools, including generation of profile plots. For genome browser visualization, normalized BigWig files were generated using deepTools bamCoverage with specified bin size, normalization method, genome size, and processing parameters.

### Statistical Analysis

Data are presented as mean ± SEM. Statistical significance was determined using two-tailed Student’s t-test for two-group comparisons or one-way ANOVA with Tukey’s post-hoc test for multiple comparisons. P<0.05 was considered statistically significant.

## Data availability

All data analyzed in the paper are available through the figshare. All other data that support the findings of this study are available upon reasonable request from the corresponding authors. All code developed for and utilized in this study are available on GitHub at this address: https://github.com/iamzhangxiaoyu. Any additional information required to reanalyze the data reported in this paper is available from the lead contact upon request. Data for RNA-seq (THY1 + SG, PS, RS) and RNA-seq (KIT + SG) were derived from GSE55060 and GSE89502, respectively. The scRNA-seq data was derived from GSE211390. Source data are provided with this paper.

## Acknowledgments

We thank Clark Bloomer, Rosanne Skinner, Veronica Cloud, and Yafen Niu at the University of Kansas Medical Center Genomics Core, which is supported by Kansas Intellectual and Developmental Disabilities Research Center (NIH U54 HD 090216), the Molecular Regulation of Cell Development and Differentiation—COBRE (P30 GM122731-03)—the NIH S10 High-End Instrumentation Grant (NIH S10OD021743) and the Frontiers Clinical and Translational Science Institute grant (UL1TR002366).

This work was supported by the NIH grants, R01HD103888 and R01GM157716, and the Department of Cell Biology and Physiology to N.W. X.Z. was a recipient of University of Kansas Medical Center Research Institute Lied Pilot Award. X.Z. was a recipient of Kansas IDeA Network of Biomedical Research Excellence Developmental Research Project Program Award from the National Institute of General Medical Sciences of the NIH under grant number P20 GM103418. The content is solely the responsibility of the authors and does not necessarily represent the official views of the above-mentioned funders.

## Author contributions

X.Z. and N.W. designed research; X.Z. and Y.L. performed research; X.Z. and N.W. analyzed data; and X.Z. and N.W. wrote the paper.

---

**Supplementary Figure 1.**
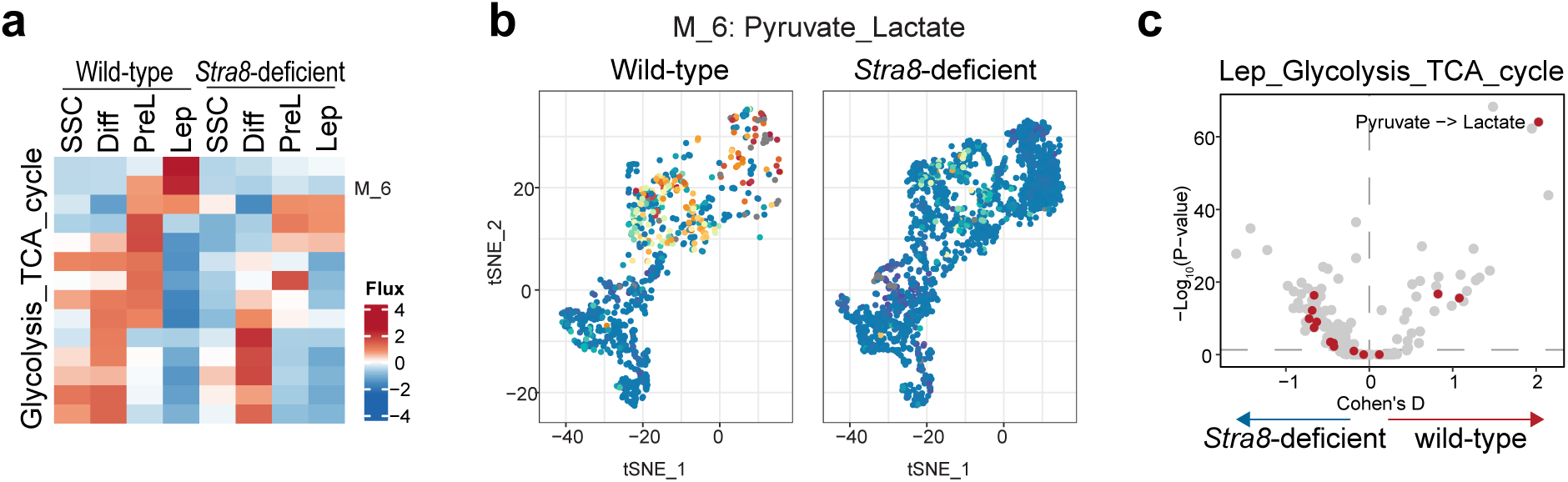
*Stra8* deficiency alters metabolic reprogramming in testicular germ cells. **a**, Heatmap of predicted metabolic fluxes for glycolysis, the TCA cycle, and Pyruvate–Lactate conversion (M_6) across testicular germ cell types (SSC, Diff, PreL, Lep) in wild-type and Stra8-deficient mice. Red and blue indicate high and low flux, respectively. **b**, tSNE projection of single cells colored by Pyruvate–Lactate conversion flux (M_6). Wild-type cells exhibit distinct populations with high metabolic flux, which are largely absent in *Stra8*-deficient cells. **c**, Volcano plot showing differentially activated metabolic reactions in Leptotene (Lep) spermatocytes between wild-type and Stra8-deficient testes. Significant reactions are highlighted in red; notably, the conversion of pyruvate to lactate is significantly activated in wild-type cells (positive Cohen’s D).

**Supplementary Figure 2.**
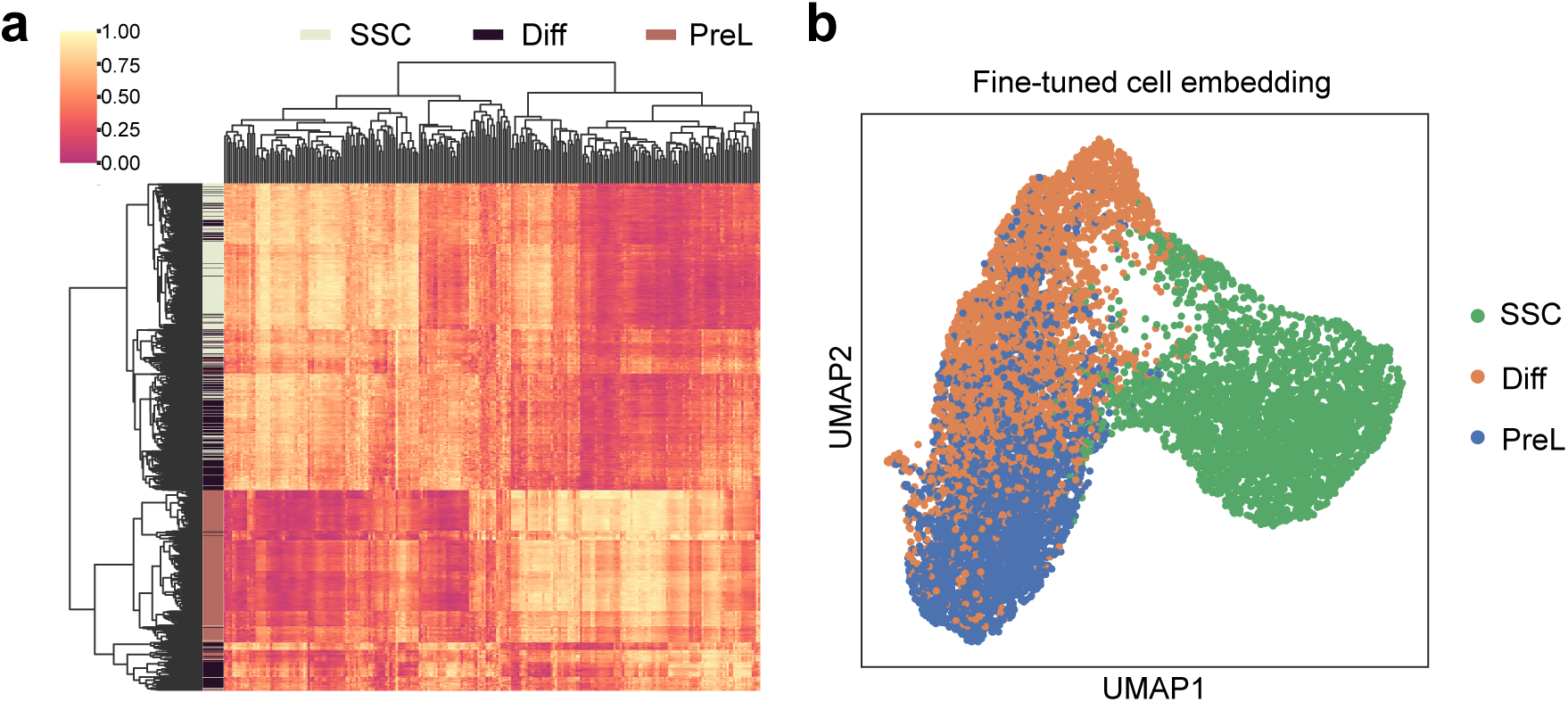
Transcriptomic profiling and fine-tuned embedding of early spermatogenic cell populations. **a**, Hierarchical clustering heatmap of gene expression profiles across three distinct cell populations: spermatogonial stem cells (SSC), differentiating spermatogonia (Diff), and preleptotene spermatocytes (PreL). The color scale represents normalized expression levels, illustrating distinct transcriptomic signatures for each cell type. **b**, UMAP visualization of fine-tuned cell embeddings, demonstrating the clear transcriptomic separation and developmental trajectories of SSCs (green), Diff (orange), and PreL (blue) cells.

**Supplementary Figure 3.**
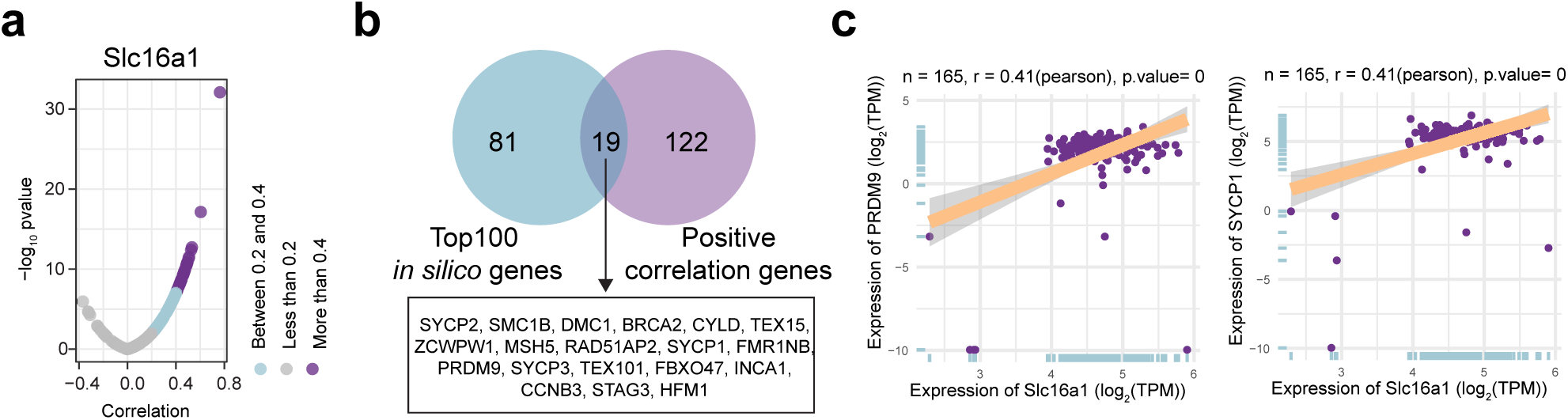
Identification of meiotic genes co-expressed with Slc16a1. **a**, Volcano plot showing the correlation of genes co-expressed with *Slc16a1*. The x-axis represents the correlation coefficient, and the y-axis represents the significance level (-log10 *P*-value). Data points are color-coded by the strength of the correlation. **b**, Venn diagram illustrating the intersection between the top 100 in silico predicted genes and genes positively correlated with Slc16a1. The 19 overlapping genes, which are associated with meiotic processes, are listed below. **c**, Scatter plots demonstrating significant positive correlations between *Slc16a1expression* and two key meiotic markers, *PRDM9* and *SYCP1* (n = 165, r = 0.41, P = 0). Pearson correlation analysis was used.

**Supplementary Figure 4:**
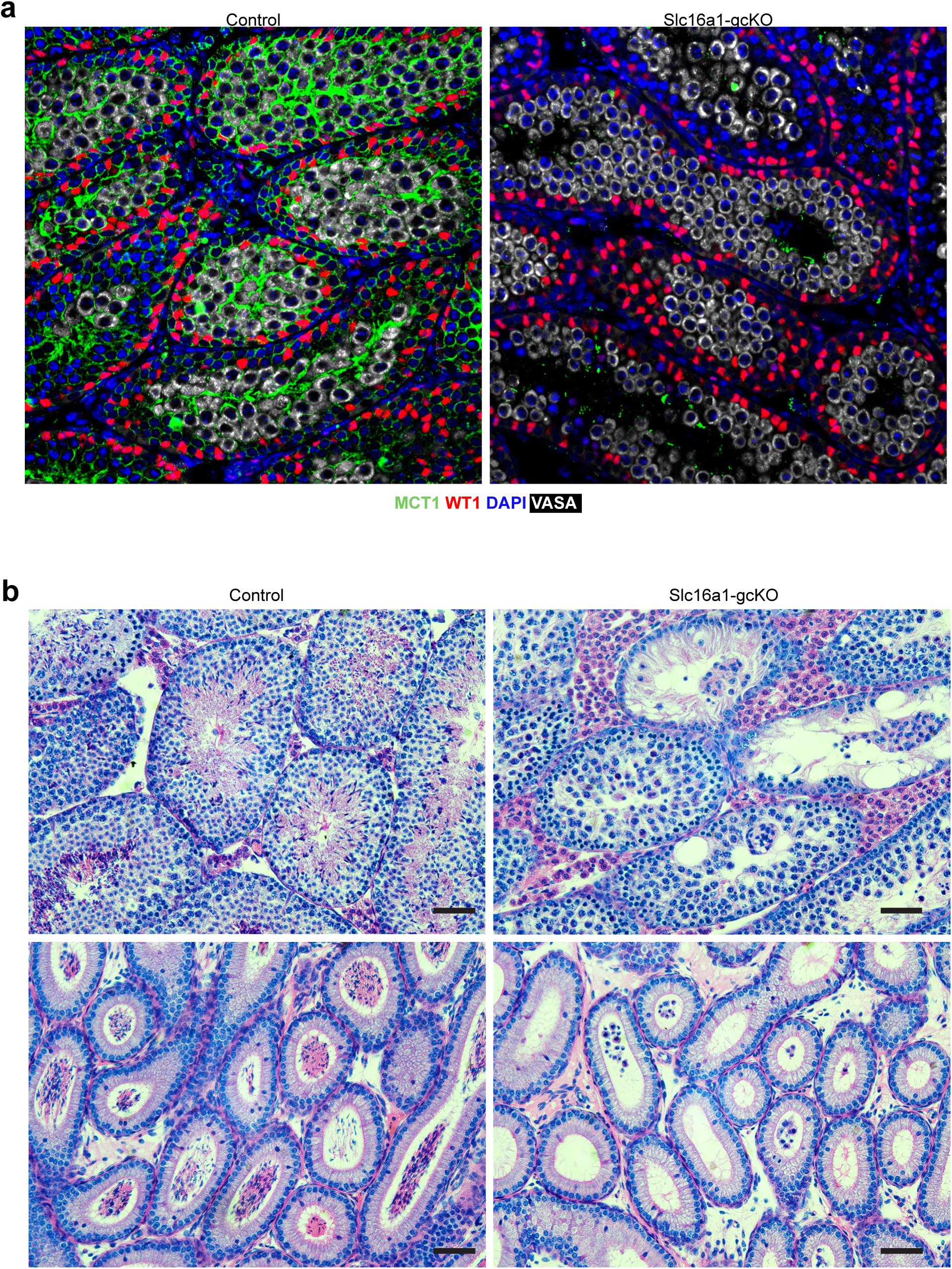
Germ cell-specific deletion of *Slc16a1* results in spermatogenic failure. **a**, Representative immunofluorescence images of Slc16a1 (green) and WT1 (red) in control and Slc16a1-gcKO testis sections. DAPI (blue) marks nuclei. The absence of Slc16a1 signal in the Slc16a1-gcKO group confirms successful conditional knockout, while WT1-positive Sertoli cells are retained. **b**, Representative H&E-stained histological sections of seminiferous tubules from control and Slc16a1-gcKO mice. Slc16a1-gcKO testes exhibit severe seminiferous tubule atrophy, extensive germ cell depletion, and vacuolization compared to the organized, full spermatogenesis observed in controls. Scale bars, 50 μm.

**Supplementary Figure 5.**
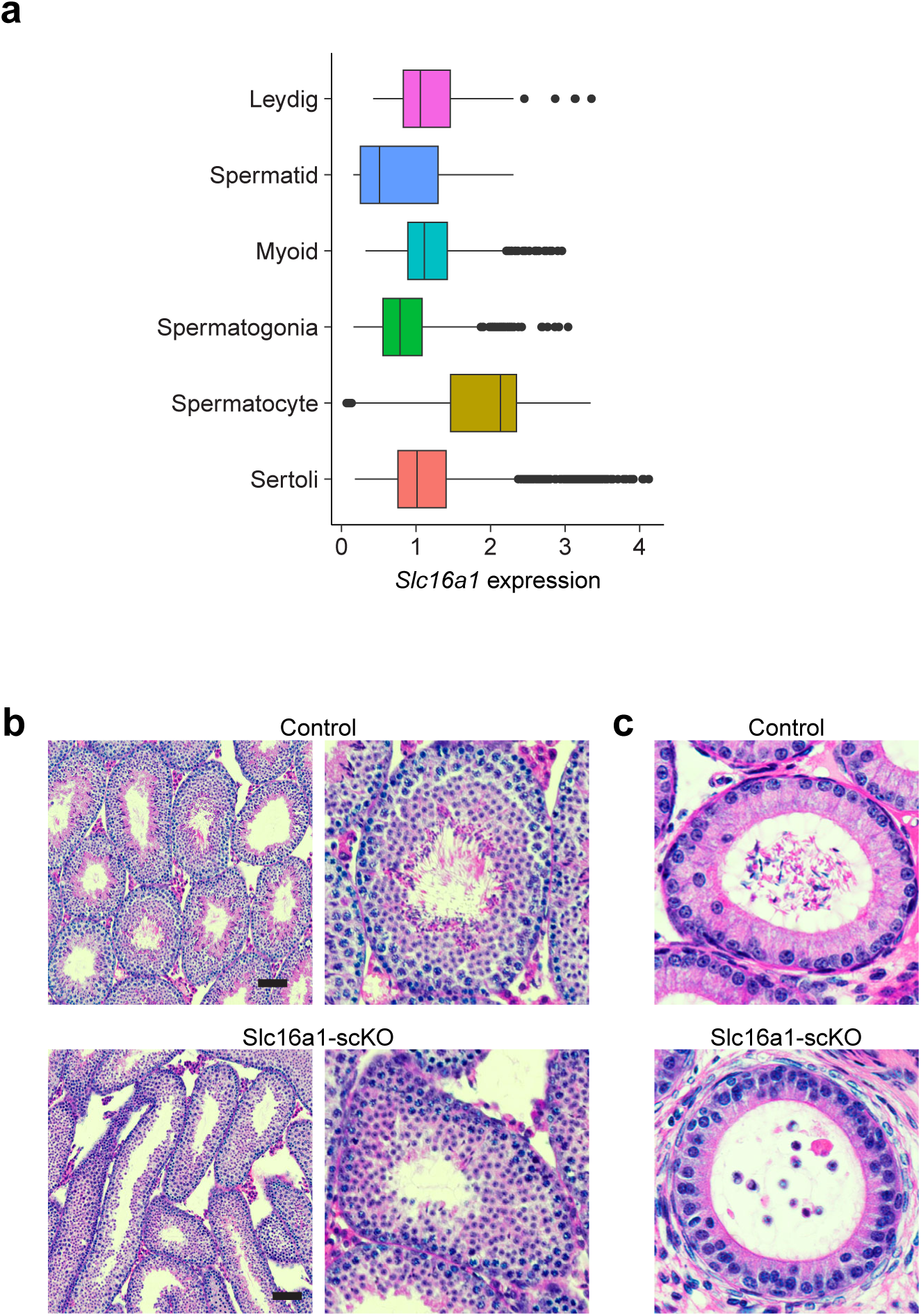
*Slc16a1* expression profile and the impact of somatic cell-specific deletion on spermatogenesis. **a**, Box plot showing the single-cell expression levels of *Slc16a1* across distinct testicular cell populations, with spermatocytes exhibiting the highest expression levels. **b**, Representative H&E-stained cross-sections of seminiferous tubules from control and Slc16a1-scKO mice. Slc16a1-scKO testes display severe tubular atrophy and a marked depletion of germ cells compared to the robust spermatogenesis observed in control testes. Scale bars, 50 μm.

**Supplementary Figure 6.**
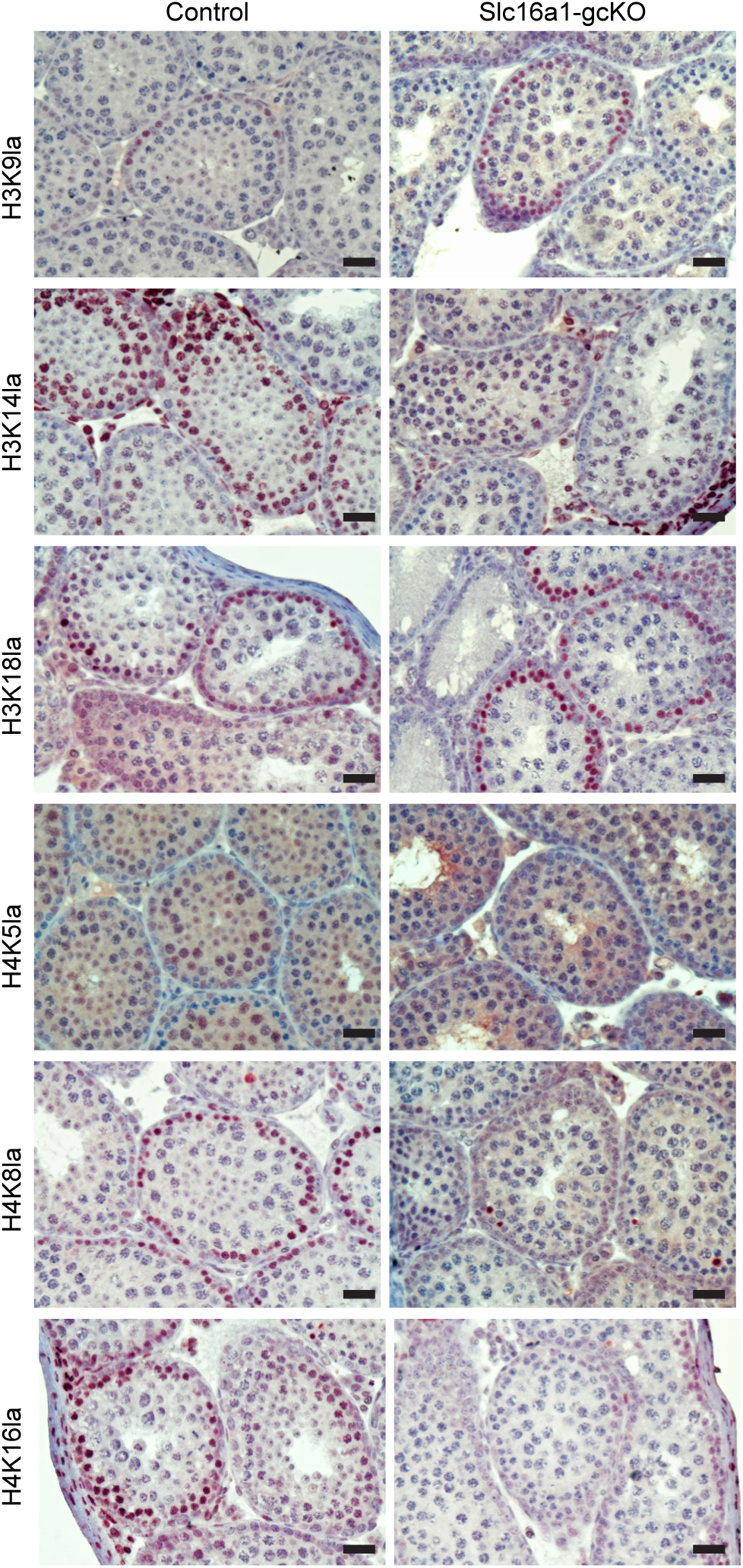
*Slc16a1* deficiency alters the histone lactylation landscape in the testis. Representative immunohistochemical staining of various histone lactylation marks (H3K9la, H3K14la, H3K18la, H4K5la, H4K8la, and H4K16la) in testicular sections from Control and Slc16a1-gcKO mice. The germ cell-specific deletion of Slc16a1 leads to differential changes in the abundance and localization of these histone marks, indicating that Slc16a1-mediated lactate transport is essential for maintaining specific histone lactylation profiles during spermatogenesis. Scale bars, 50 μm.

## Reference

1 Oulad-Abdelghani, M. et al. Characterization of a premeiotic germ cell-specific cytoplasmic protein encoded by Stra8, a novel retinoic acid-responsive gene. J Cell Biol 135, 469–477 (1996).

2 Baltus, A. E. et al. In germ cells of mouse embryonic ovaries, the decision to enter meiosis precedes premeiotic DNA replication. Nat Genet 38, 1430–1434 (2006). 10.1038/ng1919

3 Anderson, E. L. et al. Stra8 and its inducer, retinoic acid, regulate meiotic initiation in both spermatogenesis and oogenesis in mice. Proc Natl Acad Sci U S A 105, 14976–14980 (2008). 10.1073/pnas.0807297105

4 Ishiguro, K. I. et al. MEIOSIN Directs the Switch from Mitosis to Meiosis in Mammalian Germ Cells. Dev Cell 52, 429–445 e410 (2020). 10.1016/j.devcel.2020.01.010

5 Kojima, M. L., de Rooij, D. G. & Page, D. C. Amplification of a broad transcriptional program by a common factor triggers the meiotic cell cycle in mice. Elife 8 (2019). 10.7554/eLife.43738

6 Zhang, X., Gunewardena, S. & Wang, N. Nutrient restriction synergizes with retinoic acid to induce mammalian meiotic initiation in vitro. Nat Commun 12, 1758 (2021). 10.1038/s41467-021-22021-6

7 Zhang, X. et al. Transcriptional metabolic reprogramming implements meiotic fate decision in mouse testicular germ cells. Cell Rep 42, 112749 (2023). 10.1016/j.celrep.2023.112749

8 Handel, M. A. & Schimenti, J. C. Genetics of mammalian meiosis: regulation, dynamics and impact on fertility. Nat Rev Genet 11, 124–136 (2010). 10.1038/nrg2723

9 Roeder, G. S. & Bailis, J. M. The pachytene checkpoint. Trends Genet 16, 395–403 (2000). 10.1016/s0168-9525(00)02080-1

10 Soh, Y. Q. et al. A Gene Regulatory Program for Meiotic Prophase in the Fetal Ovary. PLoS Genet 11, e1005531 (2015). 10.1371/journal.pgen.1005531

11 Tu, W. B., Christofk, H. R. & Plath, K. Nutrient regulation of development and cell fate decisions. Development 150 (2023). 10.1242/dev.199961

12 Lu, V., Roy, I. J. & Teitell, M. A. Nutrients in the fate of pluripotent stem cells. Cell Metab 33, 2108–2121 (2021). 10.1016/j.cmet.2021.09.013

13 Smith, B. E. & Braun, R. E. Germ cell migration across Sertoli cell tight junctions. Science 338, 798–802 (2012). 10.1126/science.1219969

14 Boussouar, F. & Benahmed, M. Lactate and energy metabolism in male germ cells. Trends Endocrinol Metab 15, 345–350 (2004). 10.1016/j.tem.2004.07.003

15 Chen, Z. F. et al. Metabolic pathways and male fertility: exploring the role of Sertoli cells in energy homeostasis and spermatogenesis. Am J Physiol Endocrinol Metab 329, E160–E178 (2025). 10.1152/ajpendo.00074.2025

16 Jutte, N. H. et al. Regulation of survival of rat pachytene spermatocytes by lactate supply from Sertoli cells. J Reprod Fertil 65, 431–438 (1982). 10.1530/jrf.0.0650431

17 Zhang, D. et al. Metabolic regulation of gene expression by histone lactylation. Nature 574, 575–580 (2019). 10.1038/s41586-019-1678-1

18 Zhang, X., Liu, Y. & Wang, N. Dynamic changes in histone lysine lactylation during meiosis prophase I in mouse spermatogenesis. Proc Natl Acad Sci U S A 122, e2418693122 (2025). 10.1073/pnas.2418693122

19 Theodoris, C. V. et al. Transfer learning enables predictions in network biology. Nature 618, 616–624 (2023). 10.1038/s41586-023-06139-9

20 Suzuki, A. et al. Loss of MAX results in meiotic entry in mouse embryonic and germline stem cells. Nature communications 7, 11056 (2016). 10.1038/ncomms11056

21 Consortium, G. T. The Genotype-Tissue Expression (GTEx) project. Nat Genet 45, 580–585 (2013). 10.1038/ng.2653

22 Hasegawa, K. et al. SCML2 establishes the male germline epigenome through regulation of histone H2A ubiquitination. Dev Cell 32, 574–588 (2015). 10.1016/j.devcel.2015.01.014

23 Maezawa, S. et al. Polycomb protein SCML2 facilitates H3K27me3 to establish bivalent domains in the male germline. Proc Natl Acad Sci U S A 115, 4957–4962 (2018). 10.1073/pnas.1804512115

24 Liu, D. et al. Cyclin A1 is required for meiosis in the male mouse. Nat Genet 20, 377–380 (1998). 10.1038/3855

25 Kimble, J. Molecular regulation of the mitosis/meiosis decision in multicellular organisms. Cold Spring Harb Perspect Biol 3, a002683 (2011). 10.1101/cshperspect.a002683

26 Lord, T. & Nixon, B. Metabolic Changes Accompanying Spermatogonial Stem Cell Differentiation. Dev Cell 52, 399–411 (2020). 10.1016/j.devcel.2020.01.014

